# Viral Diversity Influences T Cell Responses to Enteric Human Adenoviruses F40 and F41

**DOI:** 10.1101/2025.08.05.668391

**Authors:** Holly M. Craven, Jennifer P. Hoang, Rookmini Mukhopadhyay, Arnold W. Lambisia, Benjamin A. C. Krishna, Benjamin J. Ravenhill, Charles N. Agoti, Charlotte J. Houldcroft

## Abstract

**Background:** Human enteric species F adenoviruses are a leading cause of diarrhoea-associated paediatric morbidity and mortality worldwide. The cellular immune response (antigen-specific cytotoxic T cells and secreted cytokines) to human adenovirus (HAdV) infection is known to ameliorate symptoms and is critical for viral clearance. We hypothesised that the capsid proteins (hexon and penton) of HAdV-F40 and 41 (F40, F41) are evolving to escape cellular immune responses. Major histocompatibility complex (MHC) binding of viral peptides is a key step in the presentation of peptide-MHC complexes which activate the T cell receptor and the cytotoxic T cell response.

**Methods:** Using global HAdV genomic data, we predicted MHC-peptide binding within the hexon and penton proteins of F40 and F41. We focused on MHC class I alleles common in the UK and Kenya and identifying predicted CD8^+^ T cell epitopes. Eight predicted epitope pairs from the F41 hexon were synthesised as 15mer peptides, comparing the wildtype (1970 F41 reference) to the variant (2019-2022) sequences. Cellular IFNγ responses to these epitopes were measured in healthy donors using FluoroSpot assays.

**Results:** We identified multiple predicted CD8^+^ T cell epitopes shared between HAdV-species C and F, but also unique to species F, and epitopes unique to each genotype. We show that IFNγ and IL2 peripheral blood mononuclear cell (PBMC) responses to HAdV-F are ubiquitous among healthy adult donors from Cambridge, UK. Among predicted CD8^+^ epitopes within the F41 hexon, 11/16 peptides elicited donor positive IFNγ responses from healthy donor PBMC (at least one epitope from seven out of eight peptide pairs).

**Conclusions:** The hexon and penton proteins of HAdV-F-40 and F41 are predicted to contain a number of genotype-specific, but conserved, CD8^+^ T cell epitopes which could be used to inform future vaccine design. Using the hexon of F41 as a case study, we show that predicted T cell epitopes in emergent strains are able to elicit an inflammatory cytokine response from healthy donor PBMC. The role of T cell recognition in driving enteric adenovirus evolution deserves further consideration.

## Introduction

Adenovirus is a leading cause of illness, hospitalisations and deaths due to diarrhoea in children under five worldwide (Cohen et al., 2022; Guga et al., 2022). Human Mastadenoviruses are a diverse genus of non-enveloped, double-stranded DNA viruses, of which more than 100 genotypes in seven species (grouped A-G), have been identified. The global burden of acute gastroenteritis (AGE) attributable to HAdVs is significant, with an estimated 75 million cases (Lee et al., 2020; Troeger et al., 2018). Approximately half of the cases of HAdV-associated AGE in children under five are associated with just two genotypes, the only known members of species HAdV-F: 40 and 41 (Liu et al., 2016). Over 35,000 deaths per year are attributable to F40 and F41 in children under 5 (Cohen et al., 2022). These two genotypes were initially distinguished as a result of their unique serological profiles: neutralising antibodies to F40 do not neutralise F41, and vice versa (De Jong et al., 1983). The two genotypes are 85.5% similar at a genomic level (Götting et al., 2022). In the UK, since at least the 1980s, the majority of HAdV-F AGE is caused by F41 (Cooper et al., 2000), whereas in Kenya, F40 is detected at a ratio of 1:1.2 relative to F41 (Lambisia et al., 2023). No contemporary comparable statistics exist for the UK.

Much of the HAdV genomic diversity is concentrated within the adenovirus capsid genes: hexon, penton and fibre (Lion & Wold, 2021), which are key targets for the adaptive immune response to infection (Hu et al., 2018; Sumida et al., 2005; Yu et al., 2013). It is recognised that escape from neutralising antibody responses is an important evolutionary driver in genetic diversity of species B (known to cause respiratory and urinary tract infections) and is likely also a driver of high recombination within the capsid genes of HadV-D species (known to cause ocular, gastrointestinal and respiratory diseases) (Ismail et al., 2018; Robinson et al., 2011, 2013). However, the impact of evolutionary pressure to escape from T cell recognition has been less well studied, particularly in enteric adenoviruses. For example, mice with pre-existing T cell and antibody-mediated immunity to adenovirus vectors experienced more pronounced dampening of immune stimulation by HAdV-C5 vectors than mice with passively transferred antibodies alone (Sumida et al., 2004). Additionally, the immune response to HAdV-C5 vectors was decreased by passive transfer of CD8^+^ T cells into naïve mice (Sumida et al., 2004).

Binding of viral peptides and presentation of a peptide-MHC (pMHC) class I complex is an important step in the activation of the T cell receptor (TCR) and the CD8^+^ cytotoxic T cell response (Hennecke & Wiley, 2001; Reynisson et al., 2020). Previously identified T cell epitopes for other adenovirus species have been shown to concentrate in the variable hexon and penton genes (Hutnick et al., 2010; Leen et al., 2004, 2008; Mukhopadhyay et al., 2025; Tang et al., 2006; Veltrop-Duits et al., 2006).

We and others have previously described the genetic diversity of F40 and F41 in the UK and Kenya during the periods 2015-2022 (UK) and 2013-2022 (Kenya) (Lambisia et al., 2023; Maes et al., 2023; Morfopoulou et al., 2023; Reyne et al., 2023). We hypothesised that HAdV-F40 and F41 were under selection pressure to escape from cellular immunity (antigen-specific cytotoxic T cells and secreted cytokines (Lockhart et al., 2022)) and in particular CD8^+^ T cell recognition, which has been shown to occur for viruses such as SARS-CoV-2 (Dolton et al., 2022; Stanevich et al., 2023a), EBV (Burrows et al., 1996), and influenza A (Machkovech et al., 2015).

In this study, we predicted MHC class I epitopes within the hexon and penton genes of F40 and 41 sequences from UK and Kenyan samples collected over the course of a decade (∼2013-2022), relative to the reference sequences for each genotype (isolated in the 1970s). We identified, using FluoroSpot, a group of UK blood donors with robust T cell responses to species F adenoviruses. We synthesised predicted individual variable epitopes as peptides and used these for T cell stimulation in FluoroSpot assays. This allowed us to quantify the frequency of T cells producing an interferon gamma (IFNγ) response to predicted variable epitopes. Finally, we assessed the ability of peripheral blood mononuclear cells (PBMC) derived from healthy blood donors to make an antiviral response to the diversity of adenovirus F41 hexon variants in current or recent circulation within the UK. We found that healthy blood donors could make an IFNγ response to at least one epitope from seven out of eight predicted epitope pairs.

## Materials and Methods

### Sequence Data and Alignment

Reference HAdV genome sequences were retrieved from NCBI GenBank (https://www.ncbi.nlm.nih.gov/genbank/) on January 16, 2023. DNA coding sequences (CDS) for the hexon and penton genes were downloaded from GenBank in FASTA format for reference genomes of HAdV-C1 (AC_000017.1), HAdV-C2 (AC_000007.1), HAdV-C5 (AC_000008.1), HAdV-F40 (NC_001454.1) and HAdV-F41 (DQ315364.2). DNA sequences were imported into MEGA11 (Tamura et al., 2021) and translated into protein sequences.

All F40 and F41 complete or partial genome sequences identified from metadata as originating from the UK or Kenya were retrieved from GenBank on February 7, 2023 and downloaded in FASTA format. GenBank was also searched for full-length F40 and F41 hexon and penton sequences without associated whole genome sequences, but none were identified. The DNA CDSs for the hexon and penton were extracted using MAFFT (Katoh et al., 2019) (https://mafft.cbrc.jp/alignment/server/specificregion-last.html), using default settings and ambiguous nucleotides (N) treated as a wildcard. The DNA CDSs from DQ315364.2 (strain Tak) and NC_001454.1 (strain Dugan) were used as references for F41 and F40, respectively. Extracted sequences were downloaded in FASTA format. For a full list of retrieved sequences, see Supplementary data (1. Genbank sequences).

All HAdV-F hexon and penton DNA CDSs, including reference sequences, were imported into MEGA11 and translated into protein sequences. MUSCLE in MEGA11 software with default parameters was used to generate amino acid sequence alignments for the hexon and penton. Alignments were manually corrected where necessary.

### Identification of Conserved and Variable Regions

Conserved and variable regions were identified based on previously published analyses. Hexon: (Crawford-Miksza & Schnurr, 1996); penton: (Zubieta et al., 2005).

### Identification of Common HLA Alleles

Kenyan human leukocyte antigen (HLA) frequencies were retrieved from the Allele Frequency Net Database (Gonzalez-Galarza et al., 2020) on January 19, 2023, using the “HLA classical allele frequency search” tool. Frequencies were derived from a study of Kenyan women of diverse ethnic origin. The locus was restricted to A or B. The population was restricted to “Kenya (n=144)”, with the rationale of this population being more representative of nationwide HLA allele frequencies than studies on smaller regions. All other fields were as default. HLA-A and HLA-B alleles were re-classified to two-field resolution and ranked by allele frequency. Since the catalogue of common, intermediate and well-documented HLA alleles in version 3.0.0 defines a well-documented HLA allele as having ≥5 occurrences in unrelated individuals (Hurley et al., 2020), a cut-off of 3% was chosen, which identified alleles with ≥5 occurrences in a sample size of 144 (Supplementary data 4 Kenya common HLA; 10 HLA frequencies). Alleles with frequency ≥3% were classed as “common” in Kenya.

UK HLA frequencies were obtained from two studies. One comprised individuals of Caucasian ancestry from the Oxford Biobank (n=5553) (Neville et al., 2017), chosen because of its large sample size. The second comprised individuals from the English blood donor population (n=519) (Davey et al., 2017), including those of non-British descent, chosen to partially recapitulate UK allelic variation owing to ethnic diversity. HLA-A and HLA-B alleles were re-classified to two-field resolution and ranked by allele frequency for each dataset (Supplementary data 5 UK common HLA; 10 HLA frequencies). Any allele at frequency ≥3% in either dataset was classed as “common” in the UK to allow for comparison with the Kenyan dataset.

### Prediction of CD8^+^ T cell epitopes

Peptide-MHC class I complexes which may function as CD8^+^ T cell epitopes (Hennecke & Wiley, 2001) were predicted using NetMHCpan-4.1 (Reynisson et al., 2020). Predictions were restricted to 9-mers. Strong binders (SBs) were defined by %Rank<0.5; weak binders (WBs) were defined by %Rank<2.

HAdV-C1 (AC_000017.1), HAdV-C2 (AC_000007.1) and HAdV-C5 (AC_000008.1) hexon- and penton-derived epitopes were predicted, with HLA-A and HLA-B alleles selected according to HLA restrictions of experimentally validated epitopes identified by literature review (Supplementary data 2 Published epitopes). Predictions were cross-checked with experimentally-validated epitopes (except for an HLA-B*63-restricted hexon-derived epitope, due to HLA-B*63 being unavailable in the NetMHCpan-4.1 database). For experimentally validated epitopes longer than nine residues, successful prediction was defined as prediction of any nine contiguous residues within that epitope.

C5 (AC_000008.1), F40 (NC_001454.1) and F41 (DQ315364.2) hexon- and penton-derived epitopes were predicted for HLA-A and HLA-B alleles identified as common in Kenya and/or the UK.

### Characterisation of Epitope Variation

For predicted F40 (NC_001454.1) and F41 (DQ315364.2) SB epitopes that were not shared by C5, intratypic variation was identified by manual inspection of the hexon/penton sequence alignments. Where variation was found, binding of the variant sequence(s) to HLA-A and HLA-B alleles identified as common in Kenya and/or the UK was predicted using NetMHCpan-4.1 (settings as above). The change in total allele frequency of tested HLA alleles predicted to be SBs, WBs, or NBs to the variant sequence compared to the corresponding reference sequence (NC_001454.1/DQ315364.2) was calculated in each case, using HLA frequencies from the geographic region(s) where the predicted variant epitope was identified. Both common Kenyan and UK HLA allele frequencies were considered, independent of the country-of-origin of the variant strain.

### Peptide synthesis and preparation of peptides

A custom library spanning the F41 hexon protein was synthesised by GenScript (Oxford, UK) and reconstituted into a hexon peptide pool as previously described (Mukhopadhyay et al., 2025). The F41 amino acid sequence of strain Tak (DQ315364.2) was used as the ‘wildtype’ sequence. Variable amino acid sequences predicted to contain potential T cell epitopes (see previous section) were also synthesised. These ‘variant’ sequences, listed in Supplementary data (11. Variant epitopes), were derived from Supplementary data (1. Genbank sequences) from the UK dating between 2019-2023 for eight peptides. Individual peptides were used to stimulate PBMC at a concentration of 40μg/ml/peptide.

### Ethics and donor cohort information

Ethical approval for work on samples from healthy human donors was granted for project 315068 “Understanding humoral and cellular immune responses to DNA viruses in healthy blood donors” by the HRA and Health and Care Research Wales (REC reference: 22/WA/0162). Cells were collected by the National Health Service (NHS) Blood and Transplant Service (NHSBT) in Cambridge, UK and supplied for non-clinical use.

### PBMC isolation

PBMC isolation from leukocyte reduction cones were performed as previously described (Mukhopadhyay et al., 2025). Samples were stored frozen in liquid nitrogen for batch analysis.

### Detection of Cytokine Production by FluoroSpot

Cytokine production by PBMC was quantified using a dual FluoroSpot Flex kit which was specific for human IFNγ [capture mAb (1-D1K); BAM-conjugated detection mAb (7-B6-1); Anti-BAM-490] and human IL2 [capture mAbs (MT2A91/2C95); biotinylated detection mAb (MT8G10); SA-550] (X-01A02B-10, Mabtech, Sweden).

Plates were prepared and PBMC samples thawed, treated with Benzonase Nuclease (Merck) and counted as previously described (Mukhopadhyay et al., 2025). Briefly, 1.5-2.5*10e5 cells/well of donor PBMCs in TexMACS (Miltenyi Biotech, UK) were pipetted into 96 well FluoroSpot plates, coated with either anti-human IFNγ and anti-human IL2. Peptide and protein were added in concentrations defined in Table 1, and the final volume adjusted to 150 uL/well. ImmunoCult™ Human CD3/CD28 T Cell Activator (Stemcell Technologies, UK) was used as a positive control. TexMACS media (Miltenyi Biotec, UK) with 0.01% DMSO (Sigma-Aldrich, UK) was used as the negative control. All stimulations were carried out in triplicate. Samples were left to incubate for 40 hrs before antibody detection was performed according to manufacturer’s instructions. Plates were dried in a 37°C incubator before acquisiton using an AID iSpot reader (Advanced Imaging Devices, Germany). Plates were manually quality controlled, IFNγ data were normalised and analysed as (Houldcroft et al., 2020) and IL2 data were normalised and analysed as (Krishna et al., 2024).

**Table 1.**
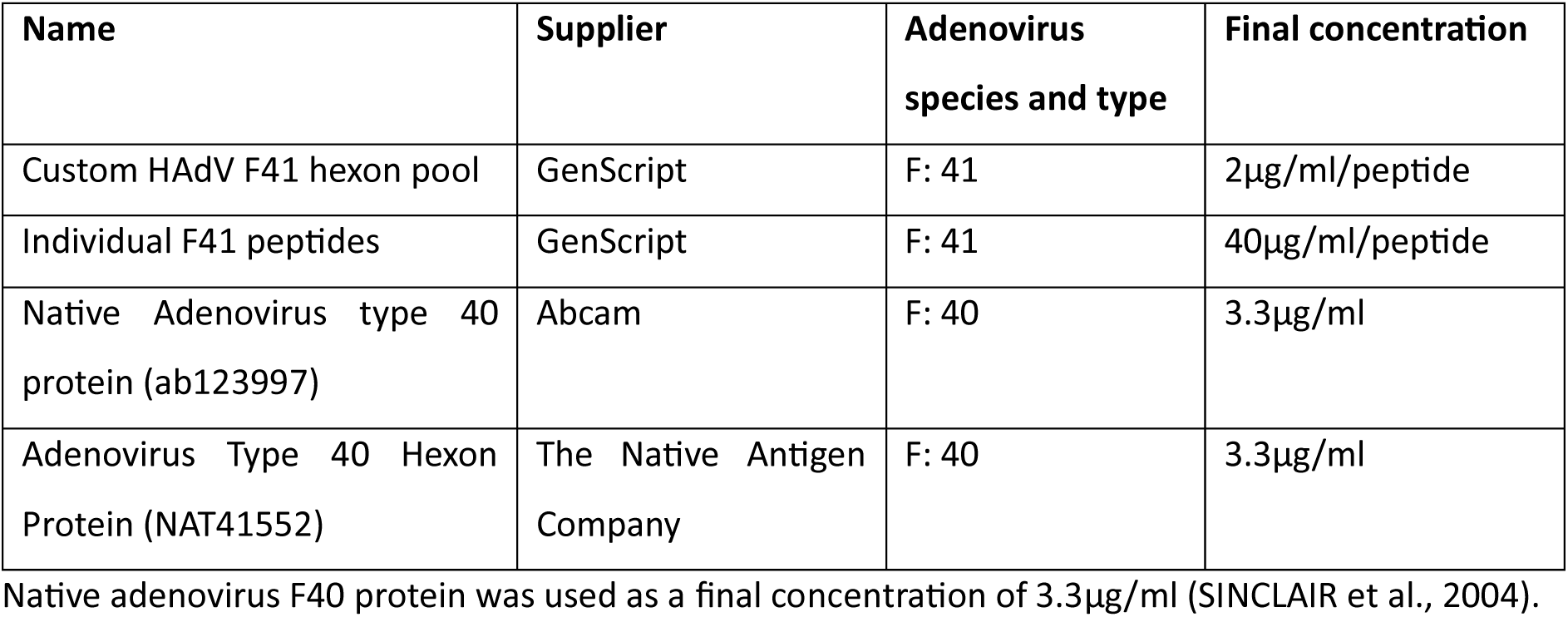

| Name | Supplier | Adenovirus species and type | Final concentration |
| --- | --- | --- | --- |
| Custom HAdV F41 hexon pool | GenScript | F: 41 | 2µg/ml/peptide |
| Individual F41 peptides | GenScript | F: 41 | 40µg/ml/peptide |
| Native Adenovirus type 40 protein (ab123997) | Abcam | F: 40 | 3.3µg/ml |
| Adenovirus Type 40 Hexon Protein (NAT41552) | The Native Antigen Company | F: 40 | 3.3µg/ml |

Native adenovirus F40 protein was used as a final concentration of 3.3µg/ml (SINCLAIR et al., 2004).

### Statistical analysis

Statistial analyses were performed and data visualised in GraphPad Prism v10.4.1.

## Results

### Identification of common Kenyan and UK HLA alleles

A single Kenyan dataset was used to identify common (defined as alleles with frequency ≥3%) Kenyan alleles, while two datasets were used for UK alleles (see methods). HLA-A and HLA-B allele frequencies were largely conserved albeit differing in frequency, and the two UK datasets were largely in accordance (Figure 1). There were eight (UK) and 11 (Kenyan) HLA-A alleles identified as common (Figure 1A), of which 4 alleles (A*01:01, A*02:01, A*03:01, A*29:02) were conserved between groups. Frequency largely varied between groups within conserved alleles however, with greater frequency of common alleles generally observed in UK datasets. The largest variation in allele frequency proportions between the UK group was observed in alleles A*01:01 (0.02 difference in frequency between the two UK datasets) and A*02:01 (0.02 difference in frequency between the two UK datasets). However, these alleles were also the two most frequent alleles in the UK datasets, whereas in Kenyan samples allele A*68:02 had greatest frequency. Kenyan group had a high frequency of ‘other’ alleles that was greater than individually observed common alleles.

**Figure 1:**
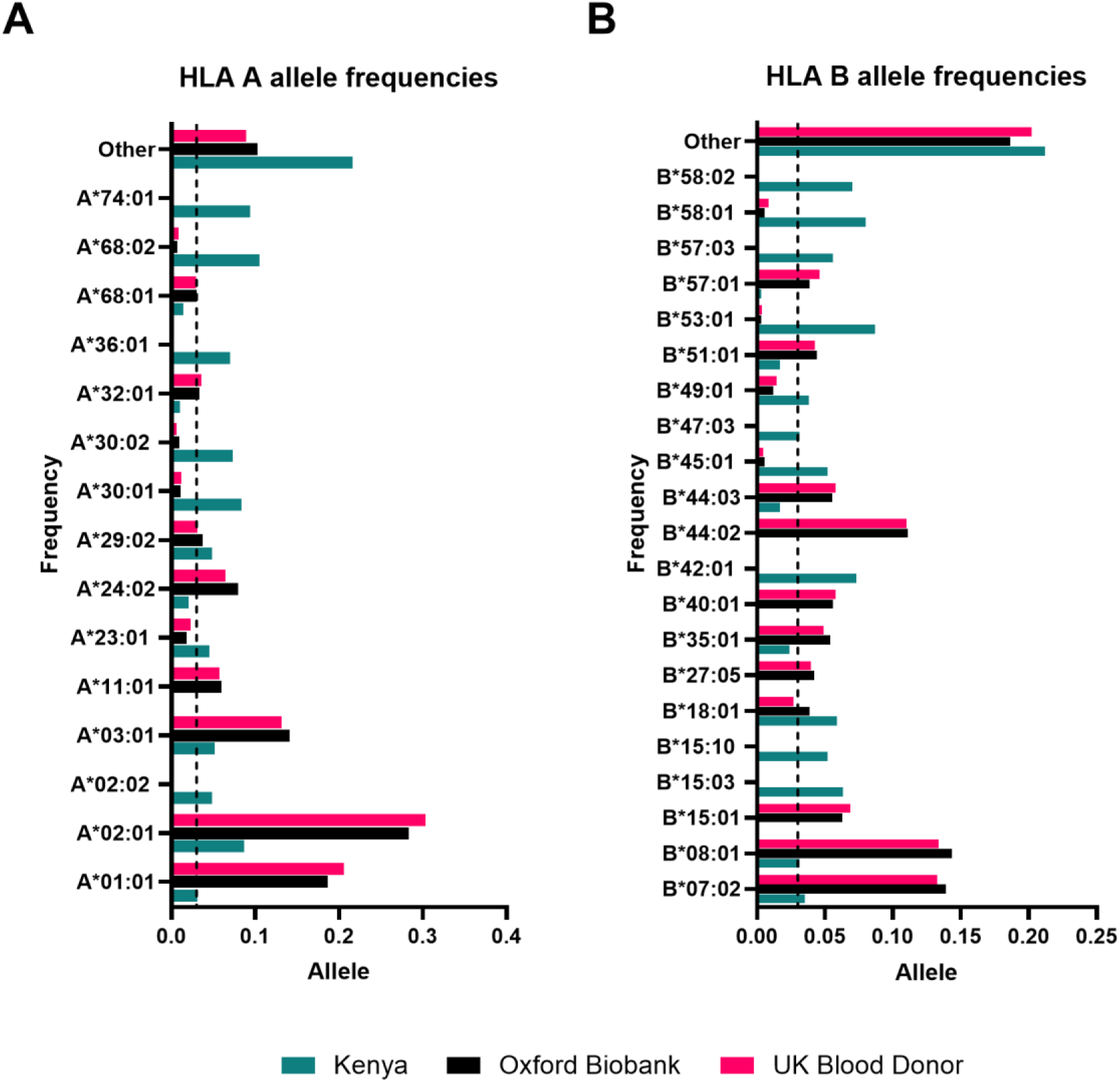
HLA allele frequency distributions for common HLA-A (A) and HLA-B (B) alleles in Kenyan and UK populations. HLA allele frequencies were obtained from UK-based studies using data from the Oxford Biobank (n=5553, shown in black) (Neville et al., 2017), a study of the UK blood donor population (n=519, shown in pink) (Davey et al., 2017); Kenyan frequencies were derived from a study of donors of diverse ethnic origin from the Allele Frequency Net Database (n=144, shown in teal) (Gonzalez-Galarza et al., 2020). HLA alleles with a frequency of ≥0.03 in one or more of these three studies are shown. Black dashed lines mark the 0.03 threshold used to define common HLA alleles (Hurley et al., 2020).

Similar patterns were observed for HLA-B alleles in both Kenyan and UK samples (figure 1B). A total of 21 alleles were recorded as ‘common’: 13 in Kenyan data and 11 in UK. Interestingly, there was a single allele that was not concordant between the two UK datasets: B*18:01. On average there was a lower frequency observed across a wider spread of alleles with greatest frequency <0.15 for both UK and Kenyan data. In all datasets, there is a large frequency (0.2) of alleles in the ‘other’ category, indicating greater spread compared to HLA-A.

Prediction of experimentally-validated HAdV CD8^+^ T cell epitopes by NetMHCpan-4.1

To evaluate the ability of NetMHCpan-4.1 to predict HAdV pMHC complexes which are CD8^+^ T cell epitopes, it was determined whether NetMHCpan-4.1 could successfully identify experimentally-validated CD8^+^ T cell epitopes of HAdV-C. From a literature review, 29 experimentally-validated CD8^+^ T cell epitopes and their respective HLA restrictions were identified for C5, 21 of which were hexon- or penton-derived (Supplementary data 2. Published epitopes). Over 70% (12/17) hexon-derived and 100% (3/3) penton-derived epitopes were identified as strong binders (SBs) or weak binders (WBs) by NetMHCpan-4.1 for C5 (Table 2). Results were consistent for genotypes C1 and C2 where epitopes were shared. The epitopes which NetMHCpan-4.1 failed to predict were not concentrated in any particular region of the hexon protein. It was concluded that, while not all epitopes could be identified, NetMHCpan-4.1 was a suitable tool for HAdV hexon- and penton-derived CD8^+^ T-cell epitope prediction.

**TABLE 2:**
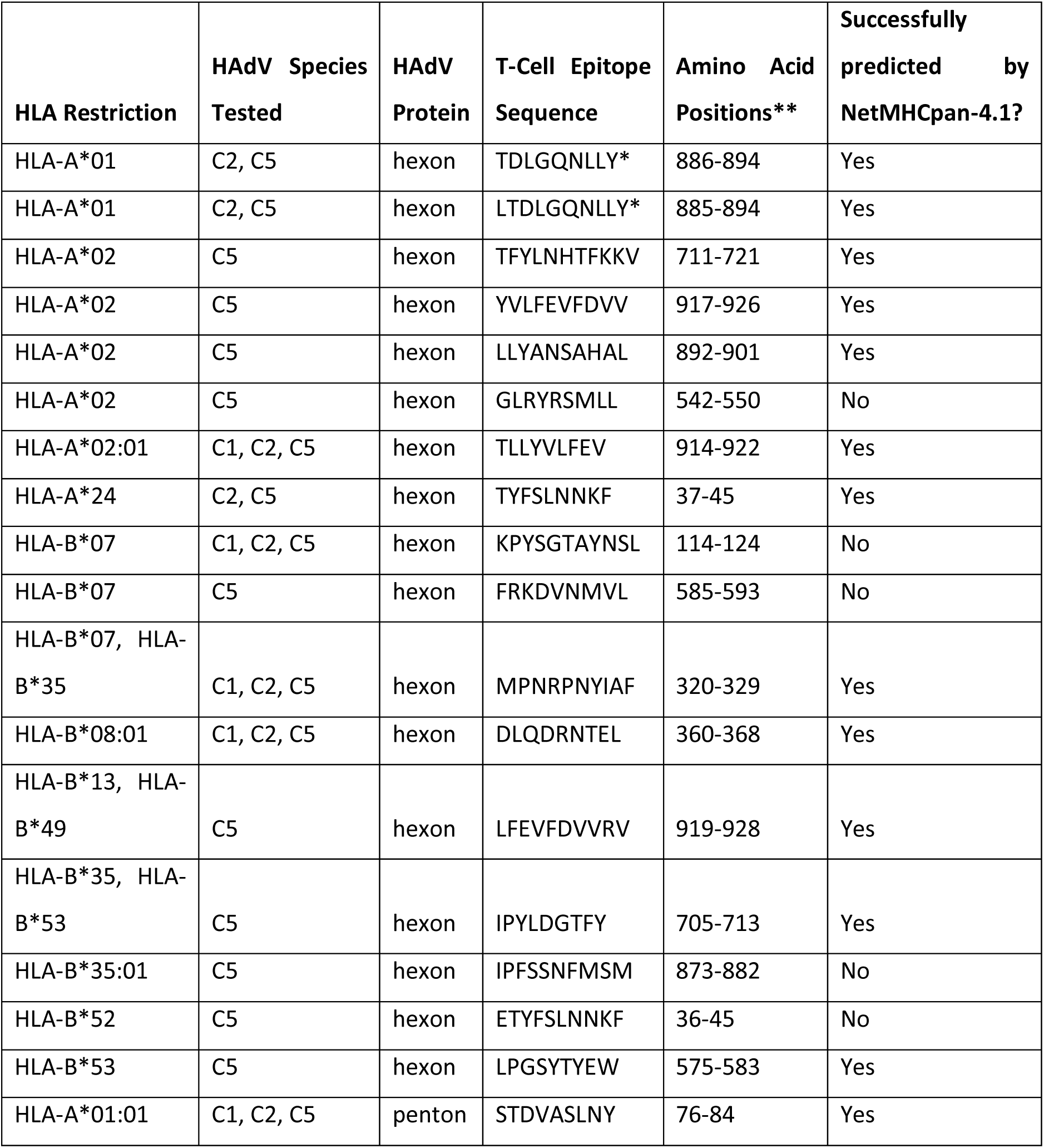
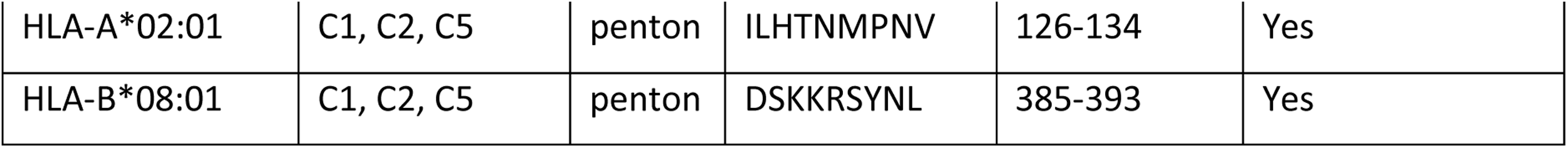
Validation of prediction of experimentally-identified T cell epitopes. *The same 9-mer was predicted for these epitopes **Amino acid positions are relative to C5 sequence

### Prediction of cross-adenovirus-species T cell epitopes

NetMHCpan4.1 was used to predict hexon- and penton-derived T cell epitopes for reference strains of HAdVs C5, F40 (Dugan) and F41 (Tak) for all common Kenyan and UK HLA alleles (see Supplementary data 4 and 5). Consistent with experimental studies, more epitopes were predicted for the hexon (supplementary data 6) than the penton (supplementary data 7). Predicted SBs were categorised as being unique to F40, unique to F41, shared by F40 and F41 only (shared HAdV-F epitopes), or shared by all three HAdVs (Supplementary data 6 - 7). No predicted SBs were unique to C5. Only shared F epitopes and unique F40/F41 epitopes were taken forward for intratypic variation analysis (supplementary data 8), since epitopes shared across HAdV species are unlikely to show intratypic variation.

### Characterisation of intratypic variation within predicted HAdV-F epitopes

To assess the intratypic variation in predicted epitopes of F40 and F41, amino acid alignments were generated using all identified hexon and penton sequences from F40 and F41 strains originating from Kenya or the UK, and those of the first sequenced F types (F41 Tak-DQ315264.2 and F40 Dugan-NC001454.1) (supplementary data 1). Mutation positions were defined relative to these alignments. UK strains spanned 2015-2022; Kenyan strains spanned 2013-2022. For F41, 38 UK hexon, 8 Kenyan hexon, 38 UK penton, and 33 Kenyan penton sequences were included. The lower number of Kenyan hexon sequences reflected missing sequence data in hexon-coding regions precluding analysis. For F40, 29 Kenyan hexon sequences and 28 Kenyan penton sequences were identified. No UK F40 sequences were available.

Phylogenetic analyses at the amino acid level (Figure 2) revealed that for the hexon (A) and penton (B), F40 and F41 formed two distinct clusters, reflecting deep intertypic divergence relative to intratypic variation (consistent with the lack of intertypic HAdV-F recombination identified by DNA-level phylogenetic analyses (Götting et al., 2022)). Within F41, hexon and penton sequences tended to cluster by country, suggestive of protein-level geographic variation. Within each type, short branch lengths indicated high conservation of the hexon and penton amino acid sequences. This included Dugan and Tak, despite their isolation in 1979 and 1973, respectively (De Jong et al., 1983).

**Figure 2:**
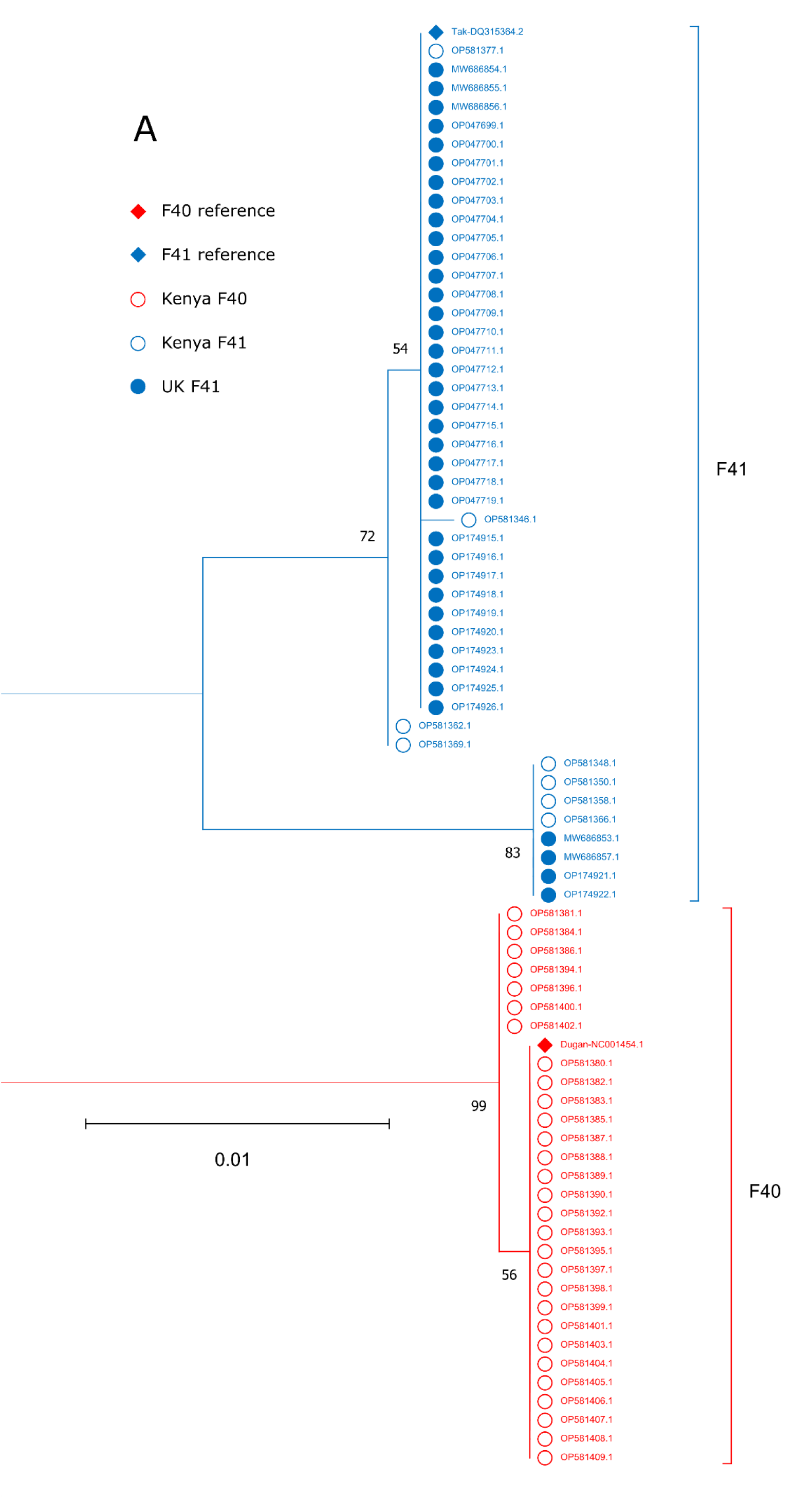
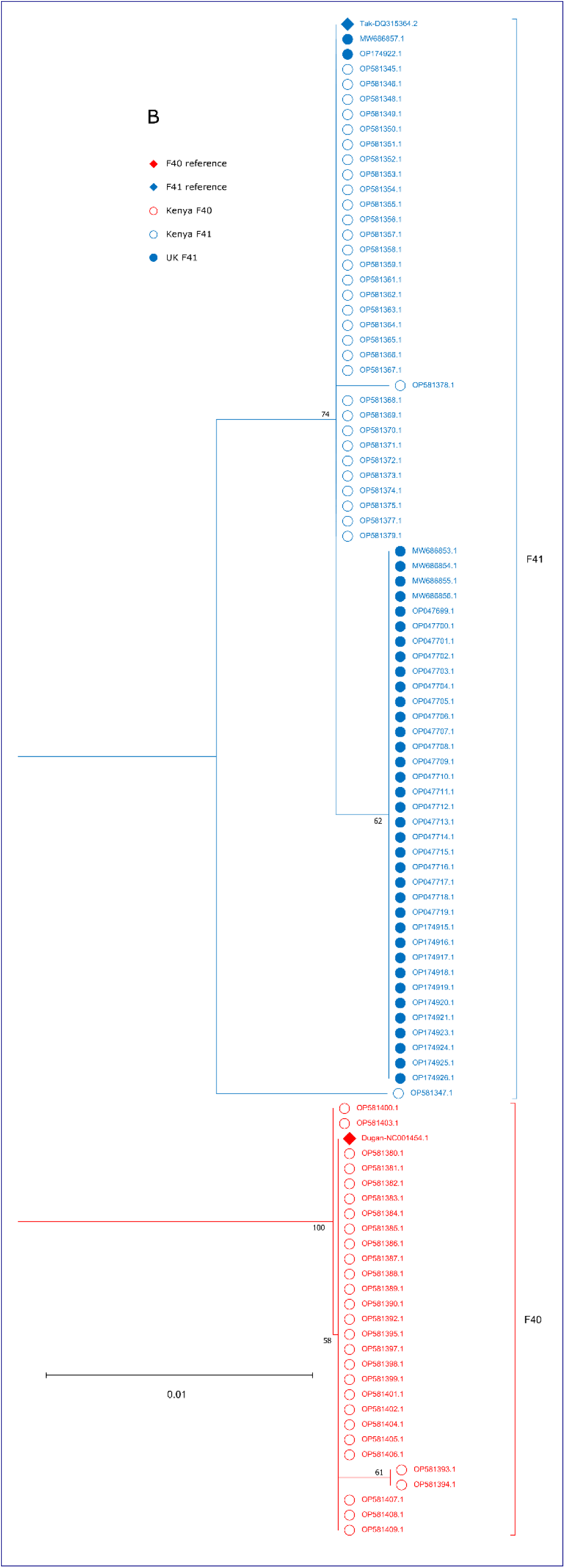
Phylogenetic trees for HAdV-F hexon (A) and penton (B) amino acid sequences. The root and total distance separating the F40 and F41 types are not shown due to the low intertypic sequence identity relative to intratypic sequence identity. F41 sequences are shown in blue; F40 sequences are shown in red. The two reference sequences (F41 Tak-DQ315264.2 and F40 Dugan-NC001454.1) are marked with a filled rhombus at the tip point followed by accession number. Sequences derived from UK samples are marked with a filled circle at the tip point followed by accession number. Sequences derived from Kenyan samples are marked with an unfilled circle at the tip point followed by accession number. Bootstrap support values are shown in black. Amino acid sequences were aligned using MUSCLE in MEGA11. Maximum likelihood trees were constructed using the amino acid substitution model WAG in MEGA11 (partial deletion, 80% site coverage cut-off).

Of the F41 unique SBs, 12 predicted hexon epitopes and 7 predicted penton epitopes showed intratypic variation (supplementary data 9). These contained only amino acid substitutions, except the hexon-derived epitope QTWTADDNY, where variant sequences displayed three amino acid substitutions (Q413G, T414N, and N421T), an amino acid insertion (insN419), or all four changes in combination. 17 F41 epitopes across the hexon and penton showed variation in UK sequences, while 9 showed variation in Kenyan sequences. However, a number of epitopes were not well covered by the Kenyan sequencing data, so there may be unsampled variation within predicted epitopes among Kenyan F41 strains. F40 appeared to show greater amino acid conservation than F41, with four predicted F40 hexon epitopes and one predicted F40 penton epitope showing intratypic variation. The four variable hexon epitopes overlapped the same single-residue substitution (A254T), while the variable penton epitope harboured a single-residue deletion in a different strain (A305del). However, there is likely unascertained F40 diversity, owing to the lack of available UK sequences and incomplete Kenyan sequencing data. No shared HAdV-F epitopes showed intratypic variation.

Mutations within predicted HAdV-F epitopes typically resided in or around previously-characterised variable regions of the hexon and penton (Figure 3). All mutations identified within predicted HAdV-F hexon epitopes resided within or near the hexon HVRs, and most predicted HAdV-F penton epitopes resided in the variable or hypervariable loops. Epitopes showing no intratypic variation (where sequence data was available for all strains) were generally located outside variable regions, though there were some exceptions (supplementary data 8).

**Figure 3:**
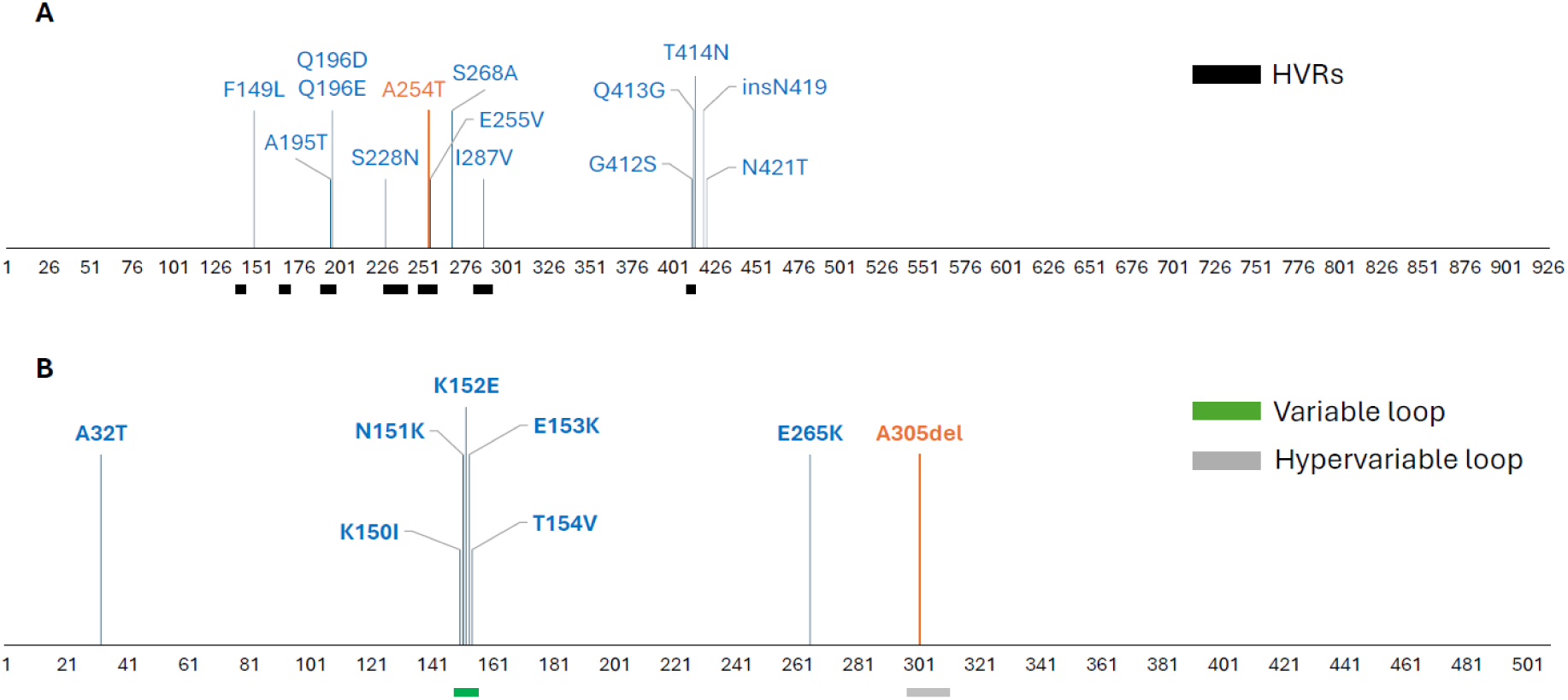
HAdV-F40 and HAdV-F41 variable epitope positions in the hexon (A) and penton (B) proteins. The horizontal axes indicate amino acid positions according to the multiple sequence alignments. Vertical lines indicate positions at which intratypic variation was identified in one or multiple predicted epitopes and are labelled with the corresponding mutation(s). F41 mutations are shown in blue; F40 mutations are shown in orange. The hexon hypervariable regions (HVRs) are indicated by black bars (left to right: HVR1 to HVR7) (Crawford-Miksza & Schnurr, 1996). The penton variable loop region is indicated by a green bar and the penton hypervariable loop region is indicated by a grey bar (Zubieta et al., 2005). Vertical line heights have been varied for clarity and do not indicate the number or type of substitution.

### Prediction of variant epitope binding mediated by common Kenyan and UK HLA alleles

To assess how intratypic variation may affect immune recognition, total allele frequencies of tested HLA alleles predicted by NetMHCpan-4.1 to be SBs, WBs or non-binders (NBs) for each variant epitope were calculated for the geographic region(s) where the variant epitope was identified. Frequencies were compared to those for corresponding reference epitopes (supplementary data 9). All identified mutations except I287V were associated with predicted binding changes for at least one epitope. In general, the total frequency of common HLA alleles predicted to be SBs decreased or remained unchanged upon mutation, though there were exceptions. The change in WB allele frequency was more variable, partly due to the counteracting contributions of SBs becoming WBs and WBs becoming NBs.

### T cell responses to HAdV-F40 lysate and F41 hexon

We have previously demonstrated that cellular immune responses to the F41 hexon are widespread and frequent in healthy blood donors (Mukhopadhyay et al., 2025). F41 is more frequently detected in clinical and wastewater samples in the UK and Ireland than in Kenya (Lambisia et al., 2023; Maes et al., 2023; Martin et al., 2023). We therefore wished to compare the cellular immune responses of the same cohort of blood donors from England to see if IFNγ and/or IL2 responses to F40 were less frequent than responses to F41.

We compared responses to the hexon protein of F41 and total F40 viral lysate, using dual-coloured FluoroSpot (SUPPLEMENTARY FIGURE 1). There was no statistically significant difference (two-way ANOVA; Tukey’s multiple comparisons test) in the frequency of PBMC IFNγ (p=0.49) or IL2 (p=0.97) responses to F40 viral lysate and F41 hexon in a cohort of 13 healthy adult blood donors from Cambridge, UK. Based on positivity thresholds from adenovirus-vectored vaccine recipients (n = 13) (Mukhopadhyay et al., 2025) 92% of donors made a positive IFNγ and 100% of donors made a positive IL2 response to F41; applying the same cutoffs to F40 viral lysate, 83% (n = 12) of donors made positive IFNγ and IL2 responses.

### T cell responses to F41 hexon variable epitopes

All donors made an IFNγ and/or IL2 PBMC response to either F40 or F41 (supplementary fig 1). However, the F40 protein and F41 hexon peptide pool are derived from HAdV-F strains Dugan and Tak, originally isolated in the 1970s (De Jong et al., 1983). Phylogenetic analysis places the F41 reference Tak in F41 lineage 1 (Götting et al., 2022), a lineage which has not been detected in the UK (Maes et al., 2023). We therefore investigated whether changes in predicted T cell epitope sites within the F41 hexon, which have accumulated between the 1970s and 2019 onwards, have led to changes in immune responses to these regions of the hexon protein. The pMHC binding predictions are based on class I HLA types, therefore we used IFNγ as a proxy for successful presentation to, and antiviral response from, CD8+ T cells (Slifka & Whitton, 2000). IFNγ responses were measured by FluoroSpot. Total PBMC were stimulated with eight pairs of 15-mer peptides, each pair containing a ‘wild type (WT)’ synthesised to match the reference sequence (1970s) and the ‘variant’ sequence synthesised based on sequences from F41 genomes derived from patients from 2019 onwards. Peptide pair locations are shown in Figure 4 and sequences in supplementary data 11.

**Figure 4:**
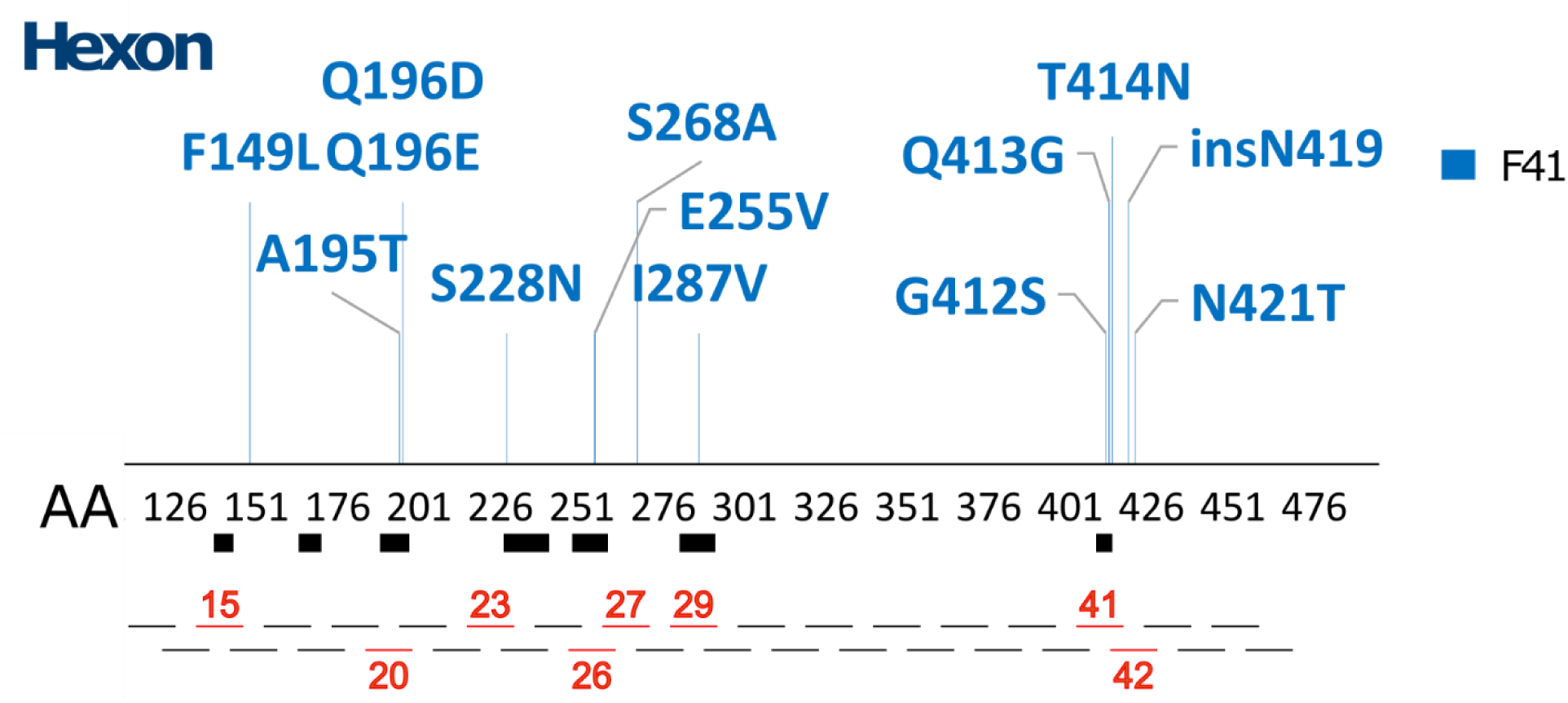
Diagram showing the location of predicted variable epitopes within the F41 hexon (truncated to amino acid positions 110-490) and the location of synthetic 15mer peptide pairs spanning these amino acid substitutions. Peptide pairs are numbered sequentially along the hexon amino acid sequence. Black boxes represent the seven hyper variable domains within the hexon protein sequence (Crawford-Miksza & Schnurr, 1996).

**Figure 5.**
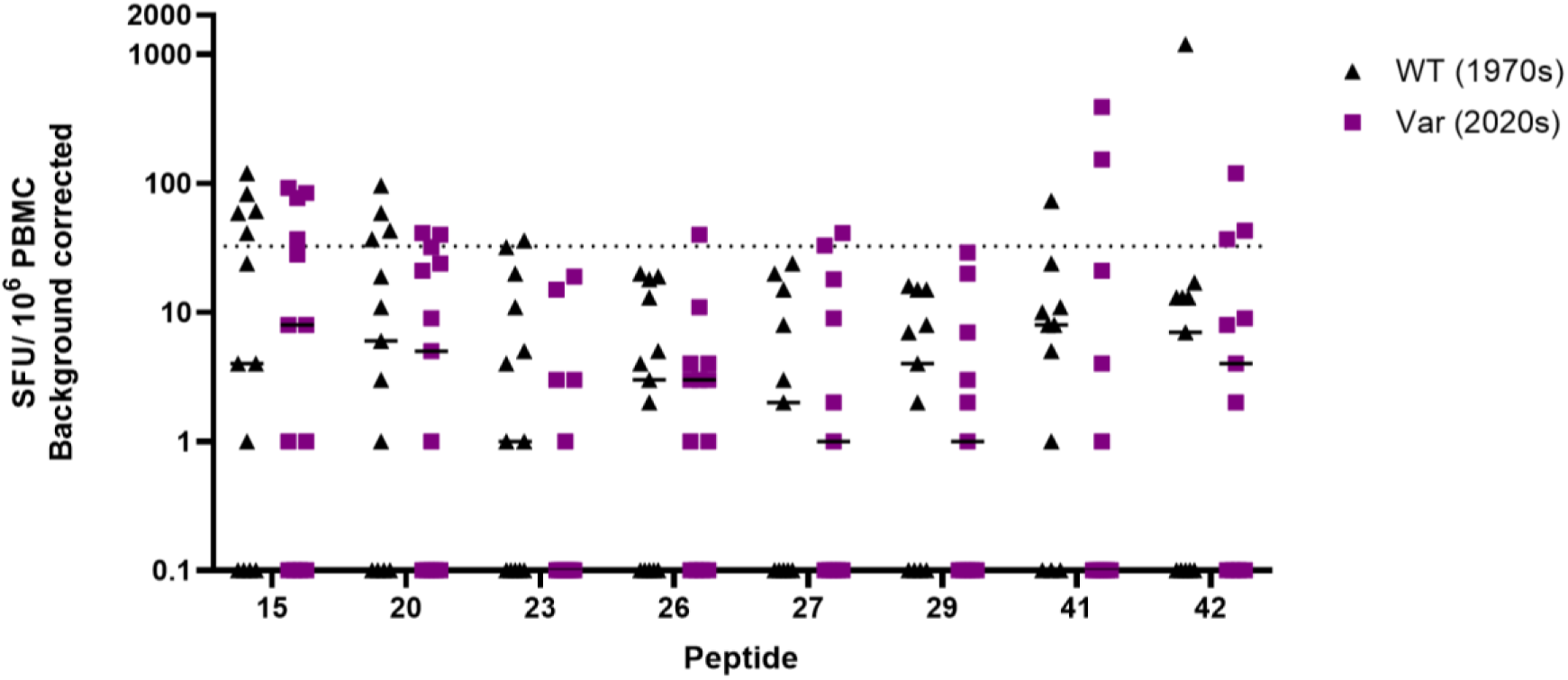
Analysis of AdV specific IFNγ FluoroSpot responses to individual peptide stimulation. Responses are calculated as spot-forming units (SFU) per 10e6 PBMC (background corrected). The dotted line indicates the boundary between positive and negative responses (Mukhopadhyay et al., 2025). WT: wild type 1970s reference sequence (black triangles); Var: 2019-2022 variant sequence (purple squares).

**Figure 6.**
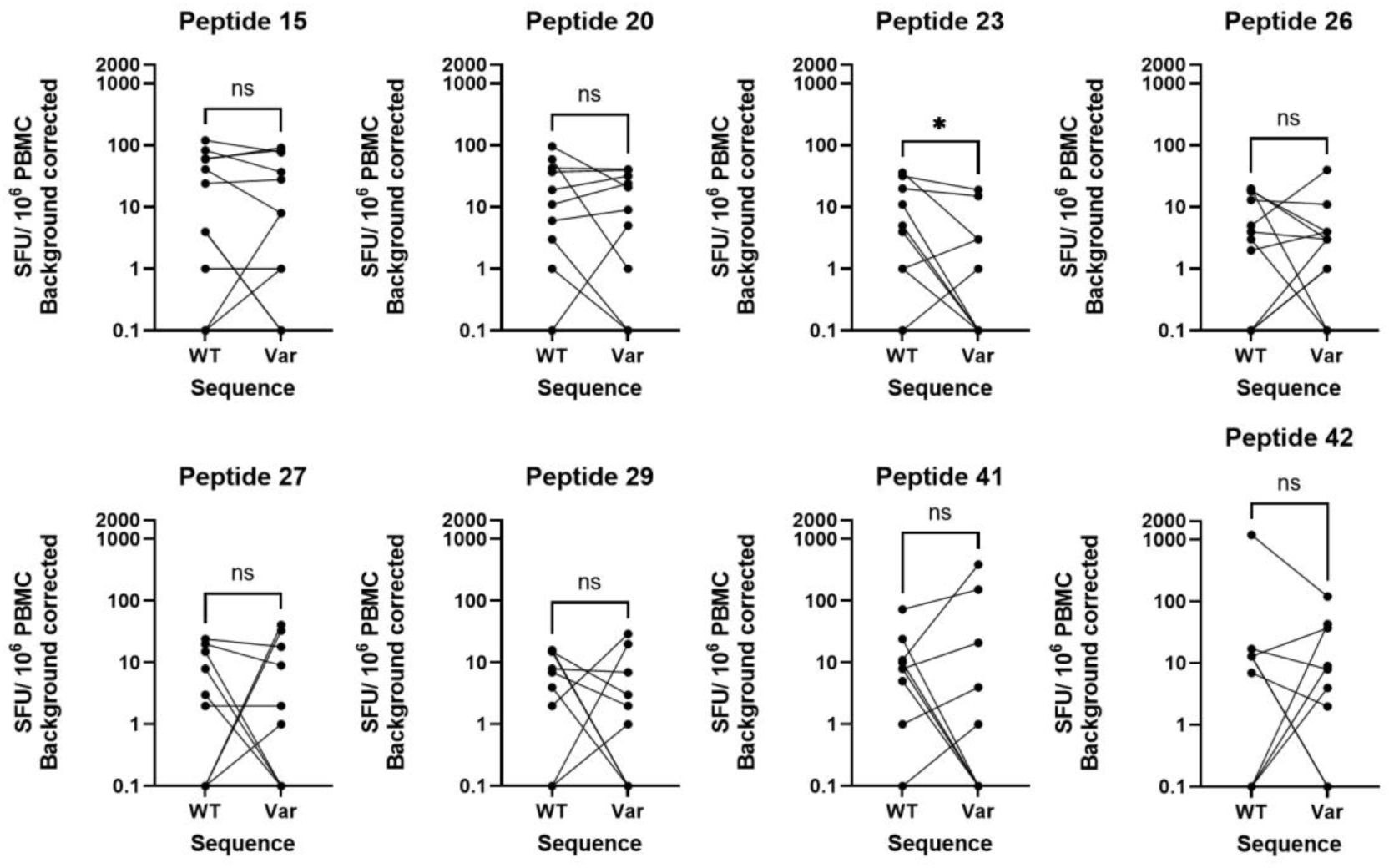
Analysis of paired within-donor PBMC IFNγ responses to F41 hexon individual peptide stimulation. PBMC from healthy blood donors were stimulated in triplicate with individual synthetic peptides representing predicted epitopes within the F40 hexon protein which have mutated between the 1970s and 2019-2022. Mean IFNγ responses were compared for each donor between pairs of peptides. WT: wild type 1970s reference sequence; Var: 2019-2022 variant sequence. Responses are calculated as spot-forming units (SFU) per 10e6 PBMC (background corrected). Significance determined by Wilcoxon matched-pairs signed rank test. Key: *p < 0.05.

The FluoroSpot results show that in seven out of eight pairs, at least one predicted CD8+ T cell epitope elicited a IFNγ response greater than 32.5 SFU in at least one donor (n = 13). The peptide to which the highest proportion of donors made a positive IFNγ response (≥32.5 SFU) was the wild type/reference for peptide 15 (38%), located between HVRs 1 and 2).

SARS-CoV-2, among other viruses, has been shown to be under evolutionary pressure to escape from both individual and population-level CD8^+^ T cell-driven immunity (Motozono et al., 2021; Stanevich et al., 2023b). We hypothesised that HAdV-F is under similar pressure, but acting on longer timescales than SARS-CoV-2. By comparing changes to each peptide within individuals, we can see that no peptide exhibits universal escape from immune recognition, nor universal gain of recognition, although mean IFNγ responses to peptide 23 were lower (p = 0.03, Wilcoxon matched-pairs signed rank test).

## Discussion

HAdV hexon HVRs have previously been identified as important in neutralising antibody escape (Roberts et al., 2006; Robinson et al., 2013). The results of this study suggest that they are also of interest in escape from CD8^+^ T cell recognition, via MHC class I binding, since mutations within predicted CD8^+^ T cell epitopes cluster around the hexon HVRs. The penton variable and hypervariable loops were similarly identified as potentially important in CD8^+^ T cell escape. For example, we predict A305del to confer partial immune escape at that epitope in a Kenyan F40 strain. Penton alignments by Zubieta and colleagues (Zubieta et al., 2005) indicate that this position coincides with the highly-conserved arginylglycylaspartic acid (RGD) motif in the penton hypervariable loop of other HAdVs. HAdV-F does not possess an RGD motif, reflecting its lack of reliance on cellular integrins for gastrointestinal cell infection (Albinsson & Kidd, 1999), so A305del may have fewer fitness effects in HAdV-F than mutations at this position in other HAdVs. This highlights how the unusual properties of HAdV-F (i.e. its enteric tropism) may permit unique mechanisms for immune escape.

Phylogenetic analyses of the hexon and penton reveal high intratypic amino acid conservation between the HAdV-F40 and F41 strains circulating over the last 10 years, and Dugan and Tak reference strains despite Dugan and Tak being isolated in the 1970s (De Jong et al., 1983). This suggests either a relatively slow mutation rate (as expected in dsDNA viruses) or strong functional constraints on the conserved regions of these proteins. This study has identified at least one predicted hexon- or penton-derived SB epitope conserved between C5, F40 and F41 for all common Kenyan and UK HLA alleles (supplementary data 6), suggesting CD8^+^ T cell epitopes can be shared across multiple HAdV species (Gardner et al., 2024; Mukhopadhyay et al., 2025). The use of conserved sequences may facilitate the development of a pan-HAdV vaccine which induces cellular immune responses (Gardner et al., 2024; Wang et al., 2023). Intratypic variation was identified in predicted unique F40 SBs and unique F41 strong binders (SBs), but not shared HAdV-F SBs, suggesting shared epitopes may reside in regions under evolutionary constraint, precluding mutations from facilitating escape from cellular immunity.

Escape from adaptive immunity is known to be an important factor in the evolution of HAdV species B and D (Ismail et al., 2018; Robinson et al., 2011, 2013), but much less is known about the evolutionary consequences of cellular immunity in driving adenovirus mutation and recombination through diversifying selection. This is true particularly within species F, where there is no evidence for intertypic recombination between the two known types (Götting et al., 2022). In the context of resurgent post COVID-19 pandemic F41 circulation (Maes et al., 2023; Reyne et al., 2023), it is important to know whether extant lineages 2A and 2B have increased immune evasiveness. This study demonstrates that computationally predicted CD8^+^ T cell epitopes can elicit antiviral IFNγ responses from healthy adult blood donors; and that while no predicted epitope was universally immune evasive in our cohort, individual regions of the F41 hexon have mutated over the last ∼50 years and these mutations have functional consequences for the magnitude of the IFNγ response within individual donors.

## Limitations to this study

MHC binding and presentation of peptides to CD8^+^ T cells is not the only factor influencing T cell response. In addition to the anchor motifs at positions 2 and 9, CD8^+^ T cell epitopes comprise TCR contact residues, which vary in position between 9-mers (Rudolph et al., 2006). TCR contact motif substitutions to analogous and heterologous sidechains do not affect MHC class I binding, but do prevent recognition by specific CD8^+^ T cells (Hsu et al., 2012). Thus, epitope mutations may alter the cellular immune response not only by precluding MHC binding, but also by affecting TCR binding to the MHC-peptide complex, a key limitation in considering only MHC-peptide binding affinity when mapping T cell epitopes. It would be instructive to characterise *in silico* how TCR binding differs between reference and variant epitopes predicted by this study. Moreover, MHC-peptide binding prediction models fail to predict all experimentally-validated T cell epitopes, thus it is possible that some amino acid variants not predicted to lie within epitope sites could, in fact, be epitope escape variants.

## Conclusions

In conclusion, candidate HAdV-F epitopes can successfully be identified *in silico* and validated *in vitro*. Some predicted epitopes are highly conserved amongst HAdVs, while others are type-specific and show intratypic variation. The results of this study illustrate the importance of jointly considering the genetic diversity of viruses and their hosts. It appears that HAdV-F is unable to mutate to escape all common HLA alleles in a given region, so vaccine candidates designed to elicit cytotoxic T cell responses may offer a promising strategy for HAdV-F control.

## Author contributions

BJR, CNA and CJH conceived, designed and supervised the study. JPH, RM, AWL, HMC, BACK and CJH performed experiments. JPH, HMC, RM, AWL, BACK, CNA and CJH performed analyses. JH and CJH wrote the original draft. All authors reviewed and edited the final work.

## Abbreviations

AGE: Acute gastroenteritis
HAdV: Human mastadenovirus
HLA: Human leukocyte antigen
NB: Non-binder
OBB: Oxford Biobank
PBMC: Peripheral Blood Mononuclear Cells
pMHC: Peptide-Major histocompatibility complex
SB: Strong binder
WB: Weak binder

